# Δ^9^-Tetrahydrocannabinolic Acid markedly alleviates liver fibrosis and inflammation in murine models of chemically- and obesity-induced liver injury

**DOI:** 10.1101/2020.05.11.088070

**Authors:** Beatriz Carmona-Hidalgo, Isabel González-Mariscal, Adela García-Martín, Francisco Ruiz-Pino, Giovanni Appendino, Manuel Tena-Sempere, Eduardo Muñoz

## Abstract

**Background:** Non-alcoholic fatty liver disease (NAFLD) is the most common liver disease in the Western world, and it is closely associated to obesity, type 2 diabetes mellitus, and dyslipidemia. Hepatocellular stellate cells (HSCs) activation by oxidative stress and inflammation is the hallmark of liver fibrosis and leads to cirrhosis and liver failure resistant to pharmacological management. Cannabinoids have been suggested as a potential therapy for liver fibrosis, prompting us to explore the antifibrotic and anti-inflammatory effects of Δ^9^-THCA-A, a major non-psychotropic cannabinoid from *Cannabis sativa* L., in animal models of NAFLD.

**Methods:** Non-alcoholic liver fibrosis was induced in mice by CCl_4_ treatment or, alternatively, by 23-week high fat diet (HFD) feeding. Δ^9^-THCA was administered daily intraperitoneally during the CCl_4_ treatment or during the last 3 weeks in HFD-fed mice. Liver fibrosis and inflammation were assessed by immunochemistry and qPCR. Blood glucose and plasma insulin, leptin and triglyceride levels were measured in HFD mice.

**Results:** Δ^9^-THCA significantly attenuated CCl_4_-induced liver fibrosis and inflammation and reduced T cell and macrophage infiltration. Mice fed HFD for 23 weeks developed severe obesity (DIO), fatty liver and marked liver fibrosis, accompanied by immune cell infiltration. Δ^9^-THCA, significantly reduced body weight and adiposity, improved glucose tolerance, and drastically attenuated DIO-induced liver fibrosis and immune cell infiltration.

**Conclusions:** Δ^9^-THCA prevents liver fibrogenesis *in vivo*, providing a rationale for additional studies on the medicinal use of this cannabinoid, as well as cannabis preparations containing it, in the treatment of liver fibrosis and the management of NAFLD.

## INTRODUCTION

Cannabis (*Cannabis sativa* L.) is a rich source of cannabinoids, a class of meroterpenoids produced in acidic carboxylated form (Hanuš et al. 2016).In countries where medicinal cannabis is legal, the plant can be assumed in a diversity of ways, from smoking and vaporizing to the consumption of cannabis-fortified baked products. All these forms of administration share a heating step. This induces the decarboxylation of acidic cannabinoids to their neutral form, as exemplified by the thermal generation of Δ^9^-tetrahydrocannabinol (Δ^9^-THC) from its precursor Δ^9^-tetrahydrocannabinolic acid (Δ^9^-THCA)(Hanuš et al. 2016).

Δ^9^-THC is the main cannabinoid from medicinal cannabis, and its biological profile underlies the use of cannabis preparations for conditions like pain, glaucoma, and the reduction of chemotherapy side effects (cachexia, vomit). On the other hand, because of its psychotropic properties, A^9^-THC also limits the medicinal uses of cannabis, even for actions conducted by other non-psychotropic bioactive constituents, including its own precursor, A^9^-THCA, which has a biological profile significantly distinct from A^9^-THC. Δ^9^-THCA is the most abundant cannabinoid from Bedrocan^®^, a cannabis variety authorized for medical use in several countries (Hazekamp et al. 2007), and cannabis tea preparations are very popular in some countries including The Netherlands and Jamaica, where decoctions are used by non-smokers and even by children and elderly people for non-recreational uses that include fever, cold, and stress (Boekhout van Solinge 1996; Janse et al. 2004). Heating associated to the preparation of cannabis teas and decoction is insufficient to decarboxylate acidic cannabinoids that, because of a much better water solubility, are selectively extracted (Hazekamp et al. 2007). Cannabis teas from dried plant material are therefore complementary to smoking or vaporizing in terms of cannabinoid profile, being enriched in acidic cannabinoids (Hazekamp et al. 2007).

Given these considerations, it is surprising that, despite their early discovery in 1955 (Krejci and Santavy 1955) acidic cannabinoids have not yet significantly benefited from the renaissance of studies on cannabis and cannabinoids. Part of this can be ascribed to the long-held assumption that acidic cannabinoids are decarboxylated in vivo and act as pro-drug of their corresponding neutral cannabinoids, a belief that has now been convincingly disproved (Crippa et al. 2020). In particular, Δ^9^-THCA is a potent activator of peroxisome proliferator-activated receptor-γ (PPARγ) (Nadal et al. 2017; Palomares et al. 2018, 2020a) and, despite not being psychotropic, can modulate CB_1_R in an orthosteric and allosteric manner (Palomares et al. 2020b) and show antiemetic activity thought CB_1_R (Rock et al. 2013).

Chronic liver disease (CLD) is a progressive degeneration of liver function that leads to liver fibrosis and cirrhosis, and eventually end-stage liver disease (ESLD) and liver failure. The principal causes for CLD are viral hepatitis, alcoholic liver disease and non-alcoholic fatty liver disease (NAFLD). Liver cirrhosis is among the leading causes for morbidity and mortality and is a risk factor for liver cancer. To date there is no cure for ESLD, and the only effective treatment is liver transplantation, which is bound to severe side effects and rely on compatible donor (Zoubek, Trautwein, and Strnad 2017). Obesity is an increasing common cause of NAFLD (Fabbrini, Sullivan, and Klein 2010; Farrell and Larter 2006) since persistent overweight leads to excessive accumulation of triglycerides into hepatocytes, *i*.*e*., steatosis, which, if maintained over time, leads to inflammation and variable degrees of liver fibrosis, eventually causing steatohepatitis (Fabbrini, Sullivan, and Klein 2010). Hence, obesity is a major risk factor for development of NAFLD (Fabbrini, Sullivan, and Klein 2010; Gholam, Kotler, and Flancbaum 2002). In turn, obesity-induced NAFLD is a main contributor to the development of major obesity-related complications, including type-2 diabetes, dyslipidemia, hypertension and cardiovascular disease (Marchesini et al. 2003), thereby exacerbating the occurrence of metabolic syndrome. Murine models, involving different nutritional manipulations, including long-term, diet-induced obesity (DIO), have been used to mimic this condition (Kristiansen et al. 2016), which defines a sort of *vicious circle* for perpetuation and aggravation of the liver and metabolic dysfunction of the obese patient. There is therefore an urgent need for effective treatments to prevent or alleviate CLD linked to obesity before its progression into a non-reversible ESLD stage. Yet, etiological treatments have largely remained elusive, and most of the pharmacological arsenal currently in use to tackle NAFLD actually targets the metabolic complications associated to this condition (Jeznach-Steinhagen et al. 2019; Musso et al. 2010). Despite less efficient, mostly symptomatic, and not devoid of adverse side-effects (Musso et al. 2010), this approach prevails due to the lack of drugs capable to efficiently block pivotal aspects of progressive liver damage, such as chronic inflammation and fibrosis.

CLD is initiated by chronic hepatocellular injury-induced oxidative stress and inflammation, resulting in the activation of hepatocellular stellate cells (HSCs) (Lee, Wallace, and Friedman 2015). This sequence of cytotoxic events can be induced by chemical agents, such as carbon tetrachloride (CCl_4_) (Dong et al. 2016; Yanguas et al. 2016). HSCs activation results in an excess of extracellular matrix deposition of alpha-smooth muscle actin (α-SMA) and collagens type I and III. Known activators of HSCs are reactive oxygen species (ROS) and immune cells-derived factors and cytokines such as platelet-derived growth factor (PDGF), transforming growth factor-beta (TGF-β), tumor necrosis factor alpha (TNF-α)and interleukin-1 (IL-1) and -6 (IL-6) delivered from immune cells, including macrophages and T cells. Reversibility of advanced liver fibrosis has been proven in patients with CLD from different etiologies, and early in the disease, cessation of causative liver injury allows fibrosis regression and activated HSCs enter apoptosis or revert into quiescent state.

Strategies for developing antifibrotic therapies aim at reducing oxidative stress and/or inhibiting-β1(Schuppan and Kim 2013). In addition, promising therapeutic targets are the CB_1_R and the PPARγ receptors. Blockade of CB_1_R and PPARγ ligand activators inhibits fibrogenesis and promotes fibrosis resolution (Jourdan et al. 2010; Zhang et al. 2013; Cinar et al. 2016). Herein, we show that Δ^9^-THCA prevents CCl_4_-induced liver fibrosis and alleviates obesity-induced NAFLD.

## MATERIAL AND METHODS

### Cannabinoids and reagents

Δ^9^-THCA-A (Δ^9^-THCA) (>97% pure) was extracted from *Cannabis sativa* L powdered flowerheads as described previously (Palomares et al. 2020b). Antibodies used in this study: anti-α-SMA (RRID: AB_262054, SC-32251, Santa Cruz Biotechnology, Dallas, TX, USA), anti-CD3 (RRID: AB_627014, SC-20047, Santa Cruz Biotechnology), anti-TNC (RRID: AB_2203818, MAB3128, R&D Systems, Minneapolis, MN, USA) and anti-F4/80 (RRID: AB_2098196, MCA497, Bio Rad, Hercules, CA, USA).

### Animals

All experiments were performed in strict accordance with European Union (EU) and governmental regulations. Handling of animals was performed in compliance with the guidelines of the European Union Directive 2010/63/EU for the use and care of experimental animals; the Ethics Committee on Animal Experimentation of the University of Córdoba (UCO, Córdoba, Spain) and the Regional Government of Andalucía approved all the procedures described in this study (2019PI/05; 03/11/14/145). Male C57BL/6 mice were used in all experiments. Measures to improve welfare assistance and monitoring of clinical status, as well as endpoint criteria, were established to minimize suffering and ensure animal welfare. Briefly, in the model of chemically-induced liver injury, involving administration of CCl_4_, wet food pellets were placed on the bed-cage when the animals began to develop clinical signs to facilitate access to food and hydration. In the model of diet-induced obesity (DIO), body weight was monitored on a weekly basis, when inspection of the animals was conducted. No overt sign of discomfort or malaise (other than excessive adiposity) was detected in HFD fed animals. Animals were housed under controlled conditions of 12 h light/dark cycles at 20°C (±2°C) and 40-50% relative humidity, with free access to water and standard food (unless otherwise state; see HFD experiments *below*).

### Induction of CCl_4_-induced liver fibrosis

Eight-week-old male C57BL/6 mice were dosed intraperitoneally (i.p.) with carbon tetrachloride (CCl_4_) (N=9-10/group), a well-known inducer of liver injury and liver fibrosis(Scholten et al. 2015), at a dose of 1 ml/kg (diluted 1:4 in corn oil, both from Sigma-Aldrich). In parallel control animals received the corresponding vehicle injections. The injections were given two times per week for 4 weeks. General health conditions, body weight and motor function of animals were periodically evaluated, twice weekly along the 4 weeks of procedure. During the 4-week CCl_4_ treatment period, mice were injected i.p. daily with vehicle (Ethanol: Cremophor: Saline 1:1:18) or Δ^9^-THCA (20 or 40 mg/kg; Supplementary information).

### Induction of high fat diet-induced liver fibrosis

Eight-week old male C57BL/6 mice were randomly assigned in two groups, and fed either a high-fat diet (HFD), D12451 (Research Diets, New Brunswick, NJ; 45%, 20%, and 35% calories from fat, protein and carbohydrate, respectively) or a control diet (CD), (A04 SAFE Diets; Augy, France; 8.4%, 19.3% and 72.4% calories from fat, protein, and carbohydrate, respectively). Nutritional intervention with HFD was maintained for a maximum period of 23 weeks to cause diet-induced obesity (DIO). Absolute body weight (BW) gain, BW gain and daily energy intake were monitored once weekly throughout the study period. The latter was calculated from mean food ingestion per week, using the energy density index provided by the manufacturer (3.34 kcal/g for CD or 4.73 kcal/g for HFD). Furthermore, the amount of fat mass and adiposity was assessed at indicated time-points during the nutritional intervention by quantitative magnetic resonance (QMR) scans, conducted using the EchoMRI™ 700 analyzer (Houston, TX, software v.2.0). Groups of mice fed with HFD were treated daily with intraperitoneal injection of Δ^9^-THCA, at a dose of 20 mg/Kg BW, for three weeks, starting from week 20 of HFD onwards. Pair-aged animals fed with HFD and treated with vehicle (Ethanol: Cremophor: Saline 1:1:18) served as controls (N= 5-6/group).

### Tissue processing

*CCl*_*4*_ *Study*: At the end of the experimental procedure, 72 h later of the last dose of CCl_4_ or vehicle, mice were euthanized, blood, liver and inguinal fat samples were collected. Tissue samples were frozen in RNA-later (Sigma-Aldrich), cooled in dry ice and stored at −80 °C for real time PCR analysis or fixed for a period of at least 24 h in fresh 4% paraformaldehyde (PFA) in 0.1M PBS (Sigma-Aldrich) for histochemical analysis. Serum samples were obtained by centrifugation (4°C, 2000 g, 20 min) within 30 min of collection and then stored at -80°C.*HFD Study*: Groups of HFD mice were euthanized at 15-, 18- and 23-weeks after initiation of the dietary intervention, when liver and inguinal fat samples were collected for monitoring progression of adipose and hepatic fibrosis. A group of male mice fed CD for 15-weeks was included for reference purposes. In addition, groups of mice fed with HFD were subjected to pharmacological intervention with Δ^9^-THCA for 3-weeks, as described above. After completion of the treatment, the animals were euthanized, and blood, liver and adipose tissue samples were collected and processed, as described above.

### Histochemical and immunohistochemistry analyses

After fixation with PFA, tissues were subjected to paraffin embedding and sectioned into 5μm-thick slices for liver and 7μm-thick slices for inguinal fat. Liver collagen was detected by PSR staining following manufacturer’s instructions (Sigma-Aldrich). For immunohistochemistry, slides were deparaffinized and rehydrated, and antigen retrieval was performed in 37 °C trypsin (pH 7.8) for 1 h or 10 mM sodium citrate buffer (pH 6) (Sigma-Aldrich) at 95 °C for 10 min. To block endogenous peroxidase activity, sections were immersed in 3.3% hydrogen peroxide in methanol (Sigma-Aldrich) for 30 min and blocked for 1 h with blocking solution (Merck-Millipore, Burlington, MS, USA) at room temperature. Tenascin (TNC; an extracellular matrix protein involved in fibrosis development), T lymphocyte and macrophage infiltration were detected with anti-TNC (1:100), anti-CD3 (1:50) or anti-F4/80 (1:50) primary antibodies overnight at 4 °C, respectively. Sections with negative controls were incubated only with PBS. Then, the slides were incubated for 1 h at room temperature with the appropriate biotin-conjugated secondary antibody: goat anti-mouse (21538, Merck-Millipore) for CD3 and goat anti-rat (BP-9400, Vector Laboratories, Burlingame, CA, USA) for TNC and F4/80. Reaction products were detected by avidin-biotin-peroxidase (Vector Laboratories) and the colour reaction was developed with DAB (3,3’ Diaminobenzidine) chromogen (Dako, Santa Clara, CA, USA) and subsequent counterstaining with haematoxylin and mounting. Imaging was performed using a light microscope Leica DM2000 microscope. The data were represented by the area percentage of each slide positive for red (PSR) or blue stain (IHC), which was calculated using Image J software (http://rsb.info.nih.gov/.ij).

### Confocal microscopy analysis

Immunofluorescent staining of α-SMA was used to identify fibrogenic myofibroblasts zones in the liver. For antigen retrieval, paraffin-embedded liver section (5 μm-thick) were deparaffinized and boiled for 10 min in sodium citrate buffer 10 mM. The sections were washed in PBS. Nonspecific antibody-binding sites were blocked for 1 h at room temperature with 3% bovine serum albumin (BSA) in PBS. Sections were then incubated with anti-α-SMA (1:500) primary antibody diluted in PBS with 3% BSA overnight at 4 °C. Sections with negative controls were incubated only with PBS. After washing in PBS, slides were incubated with goat anti-mouse Alexa 647secondary antibody (1:500, A-21235, Thermo Fisher Scientific, Waltham, MA, USA) for 1 h at room temperature in the dark. Then sections were mounted using Vectashield Mounting Medium with DAPI (Vector Laboratories). All images were acquired using a spectral confocal laser-scanning microscope LSM710, (Zeiss, Jena, Germany) with a 25×/0.8 Plan-Apochromat oil immersion lens and quantified using Image J software.

### Determination of hormonal, metabolic and inflammatory markers

Circulating levels of serum alanine aminotransferase (ALT), were measured using a commercial test kit (Sigma-Aldrich) following the manufacturer’s protocol. In addition, the circulating levels of the pro-inflammatory adipokine, leptin, were measured using the mouse leptin ELISA Kit (#90030; Crystal Chem Europe), while insulin levels were assayed using the quantitative Bio-Plex Pro™ Mouse Diabetes 8-Plex immunoassay (#171F7001M; Bio-Rad Laboratories, Hercules, CA, USA) according to the manufacturer’s instructions. Serum triglyceride levels were assayed using a GPO-POD assay kit (Triglyceride Liquid kit 992320, Química Analítica Aplicada SA, Spain).

### Intraperitoneal glucose tolerance tests

For the determination of glucose tolerance at the end of Δ^9^-THCA treatment period in HFD fed mice, animals were subjected to an intraperitoneal glucose tolerance test (iGTT). To this end, mice were ip injected with a bolus of 2 g of glucose per kg BW, after a 5 h period of food deprivation, and blood glucose levels were determined at 0, 20, 60 and 120 min after injection, using a handheld glucometer (Accu-Check Advantage®; Roche Diagnostics).

### Quantitative reverse transcriptase-PCR

Total RNA (1 µg) was isolated from liver tissues using QIAzol lysis reagent (Qiagen, Hilden, Germany) and purified with RNeasy mini kit (Qiagen) and retrotranscribed using the iScript cDNA Synthesis Kit (Bio-Rad). The cDNA was analyzed by real-time PCR using the iQTM SYBR Green Supermix (Bio-Rad) and a CFX96 Real-time PCR Detection System (Bio-Rad). Glyceraldehyde-3-Phosphate Dehydrogenase (GAPDH) gene was used to standardize mRNA expression in each sample. Gene expression was quantified using the 2^-ΔΔCt^ method and the percentage of relative expression compared to control was represented. Primer sequences are available upon request.

### Data analysis

*In vitro* data are mean ± SD and *in vivo* data are mean ± SEM. Unpaired two-tailed student T test or one-way analysis of variance (ANOVA) followed by Tukey’s post-hoc test for parametric analysis or Kruskal-Wallis post-hoc test for non-parametric analysis were used to determine the statistical significance. The level of significance was set at p □ 0.05. Statistical analyses were performed using GraphPad Prism version 8.00 (GraphPad, San Diego, CA, USA).

## RESULTS

### Effect of Δ^9^-THCA on CCl_4_-induced liver fibrosis

Eight-week-old C57BL/6 mice were randomized to healthy control group, CCl_4_-treated group in combination with vehicle (CCl_4_-vehicle), and CCl_4_-treated group in combination of 20 or 40 mg/kg of Δ^9^-THCA (Supplementary information). As reported previously, CCl_4_ induced accumulation of collagen in liver as shown by staining with picrosirius red (PSR) (Figure 1A). Δ^9^-THCA significantly reduced, in a dose-dependent manner, 23% and 31% the fibrotic liver area compared to CCl_4_-vehicle mice (Figure 1A). α-SMA content was significantly increased (8-fold) in liver from CCl_4_-vehicle compared to control mice (Figure 1B). Δ^9^-THCA significantly reduced liver α-SMA-stained area 45% and 56%, respectively, compared to CCl_4_-vehicle mice (Figure 1B). In addition, CCl_4_-induced fibrosis also increased the protein level of tenascin C (TNC), an early fibrotic marker in liver (Kasprzycka, Hammarström, and Haraldsen 2015), compared to control mice. Treatment with 20 and 40 mg/kg of Δ^9^-THCA significantly reduced liver TNC-stained area 44% and 57%, respectively, compared to CCl_4_-vehicle mice (Figure 1C). CCl_4_-induced fibrosis led to a 2.3-fold increase in ALT activity compared to control mice. High dose of Δ^9^-THCA induced a significant 1.3-fold reduction of ALT activity compared to CCl_4_-vehicle mice (Supplementary figure 2). Altogether our results showed that Δ^9^-THCA was able to prevent fibrogenesis in the liver.

**Figure 1.**
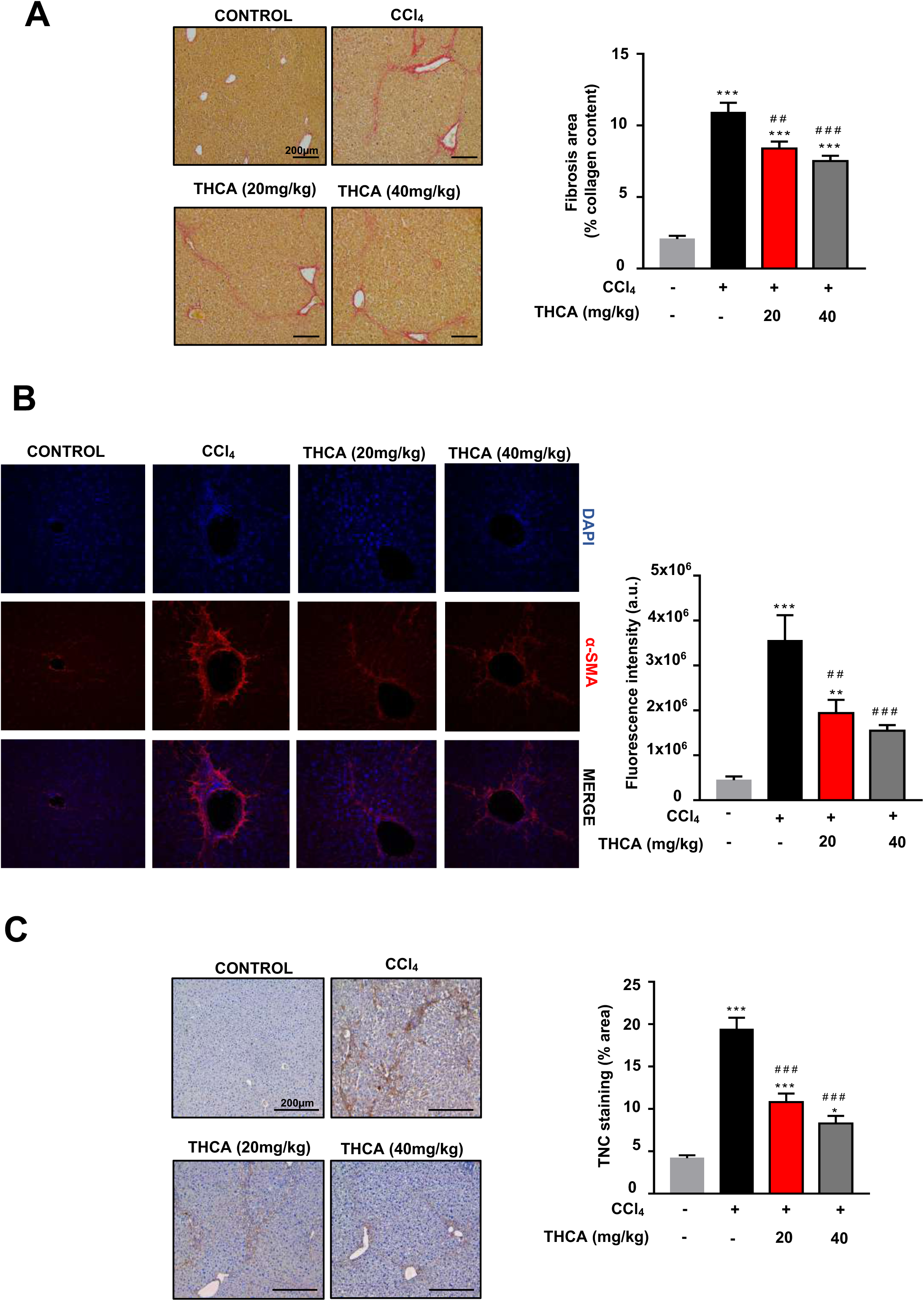
Δ^9^-THCA reduces liver fibrosis induced by CCl_4_. (A). Representative images of collagen staining in liver by picrosirius red dye (left panel) and the quantification of collagen positive area (expressed as a percentage of total liver area; right panel). (B) Representative images of α-SMA staining (red) in liver from control, CCl_4_-vehicle and CCl_4_ + Δ^9^-THCA mice is shown (original magnification x 25) (left panel). The quantification of relative α-SMA-positive area was assessed in 15-20 randomly chosen fields (right panel). (C) Representative images of immunostaining for TNC in liver (left panel) and quantification of relative TNC positive area (percentage of total liver area; right panel). Values are expressed as mean ± SEM (n=4-6 animals per group). *p< 0.05, **p < 0.01, ***p < 0.001*vs*. control group; ##p < 0.01, ###p < 0.001 *vs*. CCl_4_ group (ANOVA followed by Tukey’s test).

**Figure 2.**
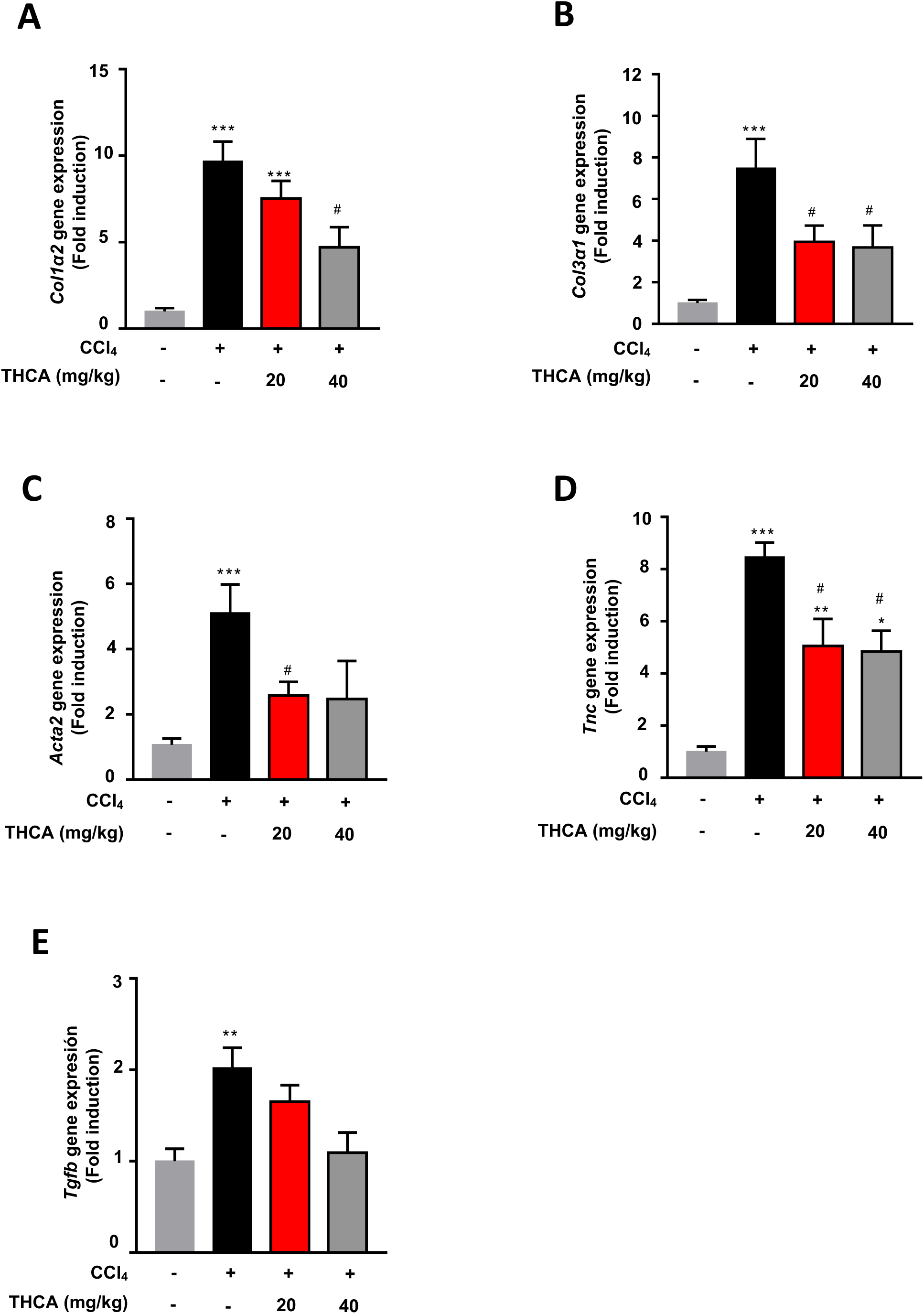
Δ^9^-THCA decreases the expression of genes involved in fibrogenesis in liver. (A) *Col1a2*,(B) *Col3a1*,(C) *Acta2* (α-SMA), (D) *Tnc* (tenascin) and (E) *Tgfb*mRNA levels in liver from control, CCl_4_-vehicle and CCl_4_ + Δ^9^-THCA mice were analysed by qPCR. Values are expressed as mean ± SEM (n=4-6 animals per group). *p< 0.05, **p < 0.01, ***p < 0.001 *vs*. control group; #p < 0.05 *vs*. CCl_4_ group(ANOVA followed by Tukey’s test).

We next analyzed the expression of genes involved in fibrogenesis in liver. As expected, CCl_4_ treatment induced a significant increase in the expression of genes involved in fibrosis compared to control mice (Figure 2A-E). High dose of Δ^9^-THCA significantly decreased the expression of *Col1a2* (Figure 2A) and *Col3a1* (Figure 2B) compared to CCl_4_-vehicle mice, while 20 mg/kg of Δ^9^-THCA only reduced the expression of the latter. Δ^9^-THCA significantly reduced the expression of *Acta2* (gene encoding for α-SMA; Figure 2C) and of the early fibrotic marker *Tnc* (gene encoding for TNC; Figure 2D) compared to CCl_4_-vehicle. However, treatment with Δ^9^-THCA had no significant effect on the expression of *Tgfb* compared to CCl_4_-vehicle, although it showed a clear tendency of inhibition (Figure 2E).

Fibrogenesis is carried out by activated HSCs upon hepatocellular injury and inflammation. Thus, we investigated the hepatic inflammation in CCl_4_-treated mice. Immunostaining for CD3 showed that CCl_4_ induces a significant 4.6-fold increase in CD3^+^ T lymphocyte infiltration into the liver compared to control mice (Figure 3A), which was reduced by 42-48% when treated with Δ^9^-THCA, independently of the dosage (Figure 3A). Similarly, liver from mice treated with CCl_4_ had 3.8-fold higher infiltration of macrophages than control mice, as seen by F4/80 staining (Figure 3B); treatment with Δ^9^-THCA induced a significant 30-38% reduction in the infiltration of F4/80^+^ cells into the liver compared to CCl_4_-vehicle mice (Figure 3B). Moreover, Δ^9^-THCA prevented in the liver the upregulation of the mRNA expression of genes coding for the inflammatory cytokines IL-1β, IL-6 and TNF-α (Figure 3C).

**Figure 3.**
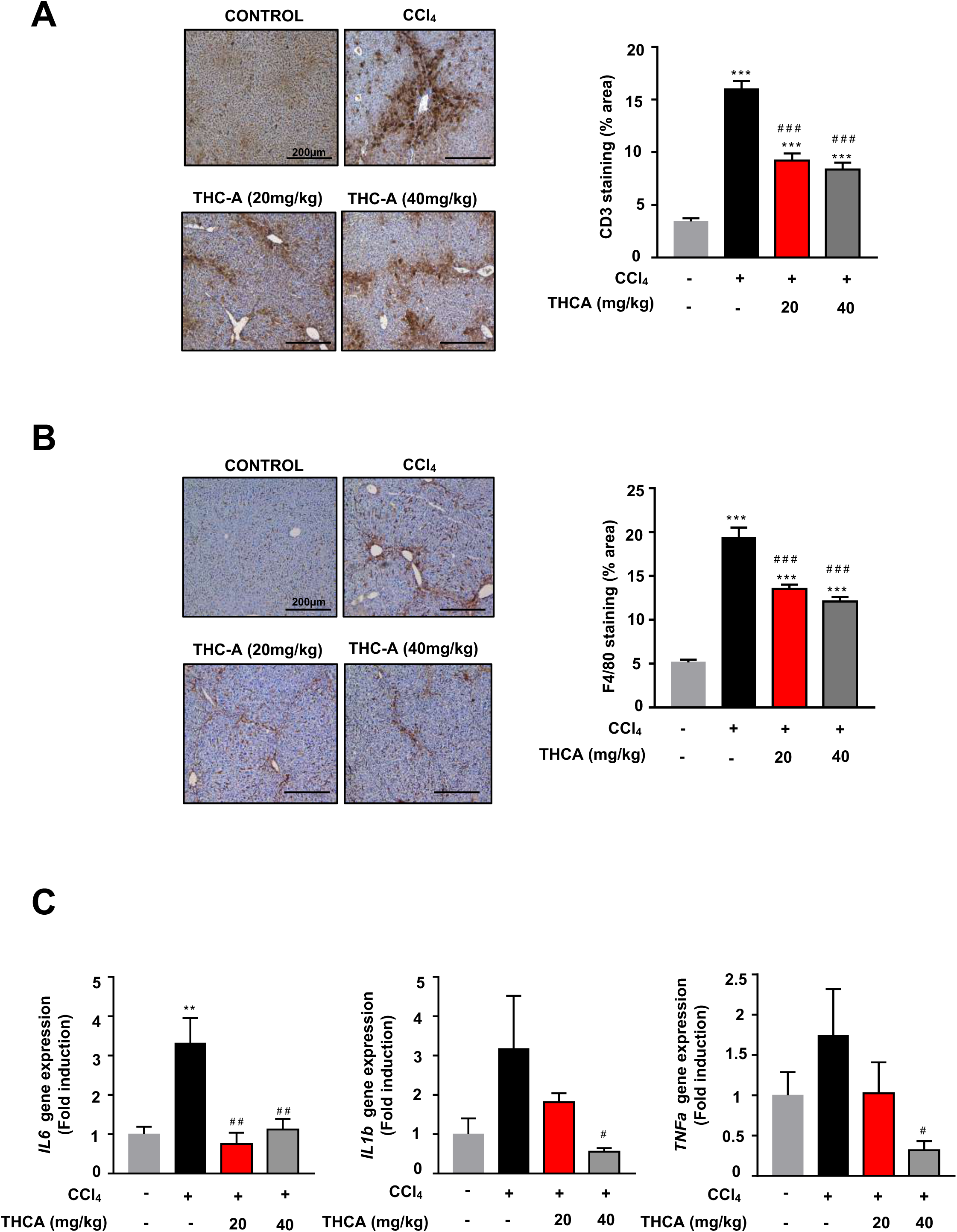
Δ^9^-THCA reduces hepatic inflammation induced by CCl_4_. (A) Representative images of immunostaining for CD3 (lymphocyte-specific marker; left panel) and the quantification of CD3^+^ cell infiltration (expressed as percentage of total liver area; right panel) in liver from control, CCl_4_-vehicle and CCl_4_ + Δ^9^-THCA mice. (B) Representative images of immunostaining for F4/80 (macrophage-specific marker; left panel) and the quantification (percentage of total liver area; right panel). (C) Δ^9^-THCA decreases the expression of pro-inflammatory genesIL-6, IL-1β, and TNF-α in theliverofCCl_4_ challenged mice. Values are expressed as mean ± SEM (n = 4-6 animals per group). **p < 0.01, ***p < 0.001 *vs*. control group; #p < 0.05, ##p < 0.01, ###p < 0.001 *vs*. CCl_4_ group (ANOVA followed by Tukey’s or Kruskal-Wallis test).

### HFD induces liver fibrosis and inflammation

The comorbidities of diet-induced obesity (DIO) include NAFLD. As fatty liver deposition progresses, inflammation increases, leading to steatohepatitis (NASH), with concomitant fibrosis. In mice, standard protocols for DIO, caused by feeding high fat diet (HFD) for 15 weeks, have been shown to induce hepatic steatosis, but failed to produce overt liver fibrosis on its own, which seems to require longer periods of HFD (Ito et al. 2007). In order to generate a model of HFD-induced liver fibrosis, we fed eight-week-old C57BL/6 male mice HFD for 23 weeks (23-wk-HFD) and analyzed the progression of liver steatosis and NASH beyond the period of 15 weeks of HFD exposure. In line with previous references(Palomares et al. 2018, 2020a), feeding HFD for 15-weeks evoked a significant increase in body weight (BW), over corresponding values in control diet (CD)-fed mice, which further increased after 18- and 23-weeks of HFD (Figure 4A). At week 15 of HFD, accumulation of triglycerides in liver was noticeable, although fibrosis was only present around the ducts and blood vessels (Figure 4B). Liver fibrosis was significantly triggered after 18 weeks of HFD, reaching 26% of liver area by 23 weeks (Figure 4B). Immune cell infiltration was present in liver tissue since week 15 of HFD (Figure 4C-D). Infiltration of CD3^+^ T cells (Figure 4C) and of F4/80^+^ macrophages (Figure 4D) was significantly higher by 15 weeks of HFD compared to CD-fed mice, and further increased by week 23 of HFD, reaching 24% and 9% of liver area, respectively. In white adipose tissue, presence of crown-like structures (CLS) (accumulation of F4/80^+^ cells) was visible after 15 weeks of HFD compared to CD-fed mice, and longer time of HFD feeding did not further increase their number (Figure 4E). These data showed that hepatic fat accumulation and liver inflammation initiates earlier than 15 weeks of HFD, while liver fibrosis occurs between 18- to 23-week-HFD in C57BL/6 male mice. Thus, our 23-week-HFD mouse model recapitulates NASH development.

**Figure 4.**
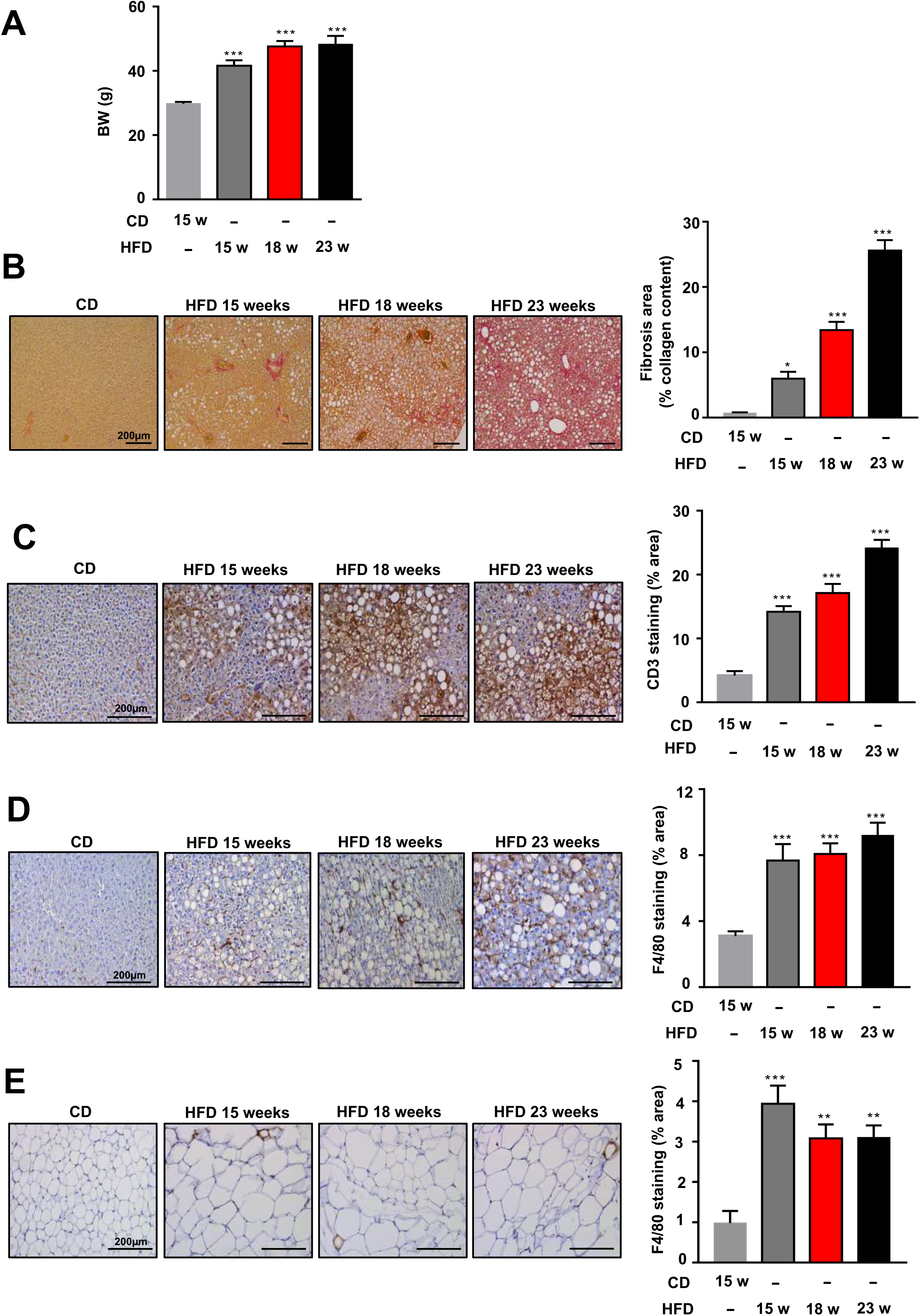
Development of a mouse model of DIO-induced NASH in mice. (A) Body weight (BW) at 15 weeks of control diet (CD) and the progression at 15, 18, 20 and 23 weeks of high fat diet (HFD) feeding in adult male mice. (B) Representative images of liver collagen staining by picrosirius red dye (left panel) and the quantification of collagen positive area (right panel) in mice fed a CD or an HFD for 15, 18 and 23 weeks. (C) Representative images of CD3-immunostained liver sections (left panel) and the quantification of relative CD3 positive area (right panel). (D) Representativeimages of immunostaining for F4/80 in liver sections (left panel) and the quantification (right panel). (E) Presence of CLS in inguinal white adipose tissue(iWAT) by immunostaining for F4/80 marker (left panel) and the corresponding quantification (right panel). Values are expressed as mean ± SEM (n=3-4animals per group). *p< 0.05, **p < 0.01, ***p < 0.001 *vs*. CD group (ANOVA followed by Tukey’s or Kruskal-Wallis test).

### Δ^9^-THCA alleviates HFD-induced liver fibrosis and inflammation, and metabolic markers

In the above model of severe, long-term obesity, during the 3 last weeks of HFD, mice were injected i.p. daily with vehicle or Δ^9^-THCA (20 mg/kg), and different metabolic and liver parameters were evaluated. Despite the massive body weight gain caused by long-term HFD, 3-wk treatment with Δ^9^-THCA caused a marked suppression of BW in obese mice, with ∼10% drop in final BW after the 3-wk treatment (Figure 5A-B). This effect was paralleled by significant changes in daily food intake, which was increased in HFD mice treated with vehicle (*vs*. CD mice), but significantly suppressed by Δ^9^-THCA treatment in obese mice (Figure 5C). Alike, in terms of body composition, HFD feeding for 23-weeks caused a marked increase in total fat mass, without changing lean mass, and enhanced the adiposity index (defined as the quotient between fat mass and the sum of fat and lean mass), while 3-week treatment of DIO mice with Δ^9^-THCA significantly reduced both parameters (Figure 5D-F). In line with previous data in less severe models of HFD-induced obesity (Palomares et al. 2020a), 23-wk feeding on HFD caused a massive glucose intolerance, as denoted by impaired GTT. Such intolerance was markedly attenuated by treatment with Δ^9^-THCA (Figure 5G-H), which lowered also the circulating triglyceride (TG) levels (Figure 5I), despite the severe obese phenotype. In good agreement, Δ^9^-THCA administration for 3 weeks to HFD mice attenuated the rise of circulating leptin and decreased serum insulin concentrations (Figure 5J-K), likely reflecting reduced insulin resistance.

**Figure 5.**
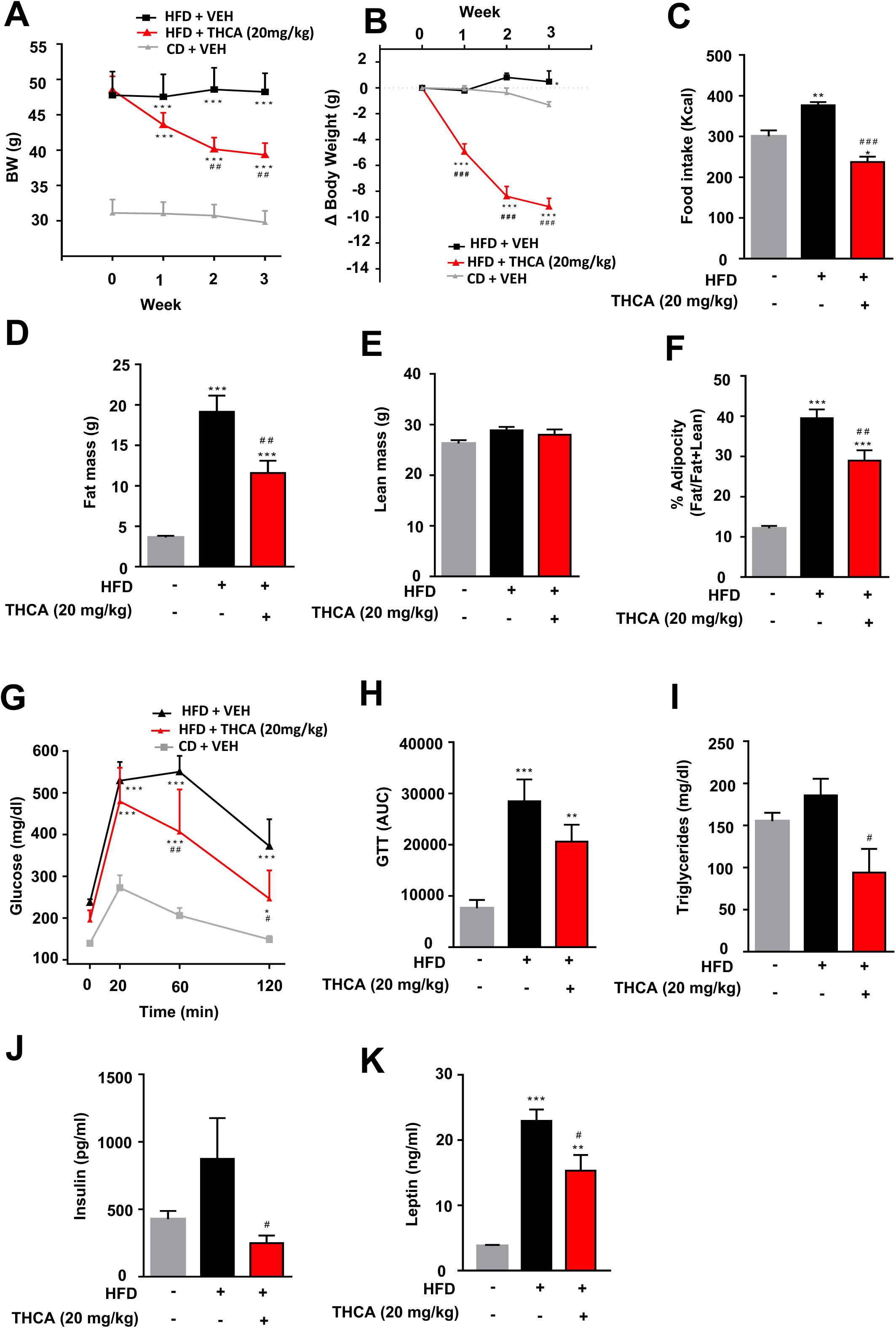
Impact of administration of Δ^9^-THCA on metabolic parameters linked to energy homeostasis in a mouse model of DIO-inducedliver fibrosis. (A, B) BW evolution of mice treated for 3 weeks with Δ^9^-THCAor vehicle after 20 weeks of HFD feeding. BW change in HFD treated mice for 3 weeks with vehicle and CD mice treated with vehicle; values are referenced to BW at the beginning of treatment (taken as 0). (C) Effect of 3-week treatment of Δ^9^-THCA on total calorie intake (Kcal) during the treatment period for the experimental groups. (D) Fat mass, (E) lean mass and (F) adiposity index, calculated as the ratio between fat mass and fat + lean mass at the end of treatments in the 3 experimental groups. (G) Glucose tolerance tests in HFD mice treated with Δ^9^-THCA or vehicle for 3 weeks and CD mice treated with vehicle and (H) integral glucose levels estimated as AUC by the trapezoidal rule in the same animals. (I) Serum triglycerides levels and plasma (J) insulin and (K) leptin levels at the end of the 3-week treatment period for the experimental groups. Values are expressed as mean ± SEM (n= 5animals per group). *p< 0.05, **p < 0.01, ***p < 0.001 *vs*. CD group; #p < 0.05, ##p < 0.01, ###p < 0.001*vs*. HFD group.

Importantly, Δ^9^-THCA significantly reduced liver fibrosis compared to 23-wk HFD-vehicle mice (Figure 6A). Livers from HFD-Δ^9^-THCA mice showed 12% fibrotic area, as assayed by PSR staining (Figure 6A), comparable to the 13% observed in 18-week-HFD mice (Figure 5B). In addition, Δ^9^-THCA significantly reduced T cell infiltration into the liver compared to 23-week HFD-vehicle mice (Figure 6B), reversing the infiltration to levels even lower than those observed in 15-week-HFD mice (Figure 4C). Similarly, Δ^9^-THCA significantly reduced the infiltration of macrophages compared to 23-week HFD-vehicle mice, to levels comparable to CD-fed mice (Figure 6C).

**Figure 6.**
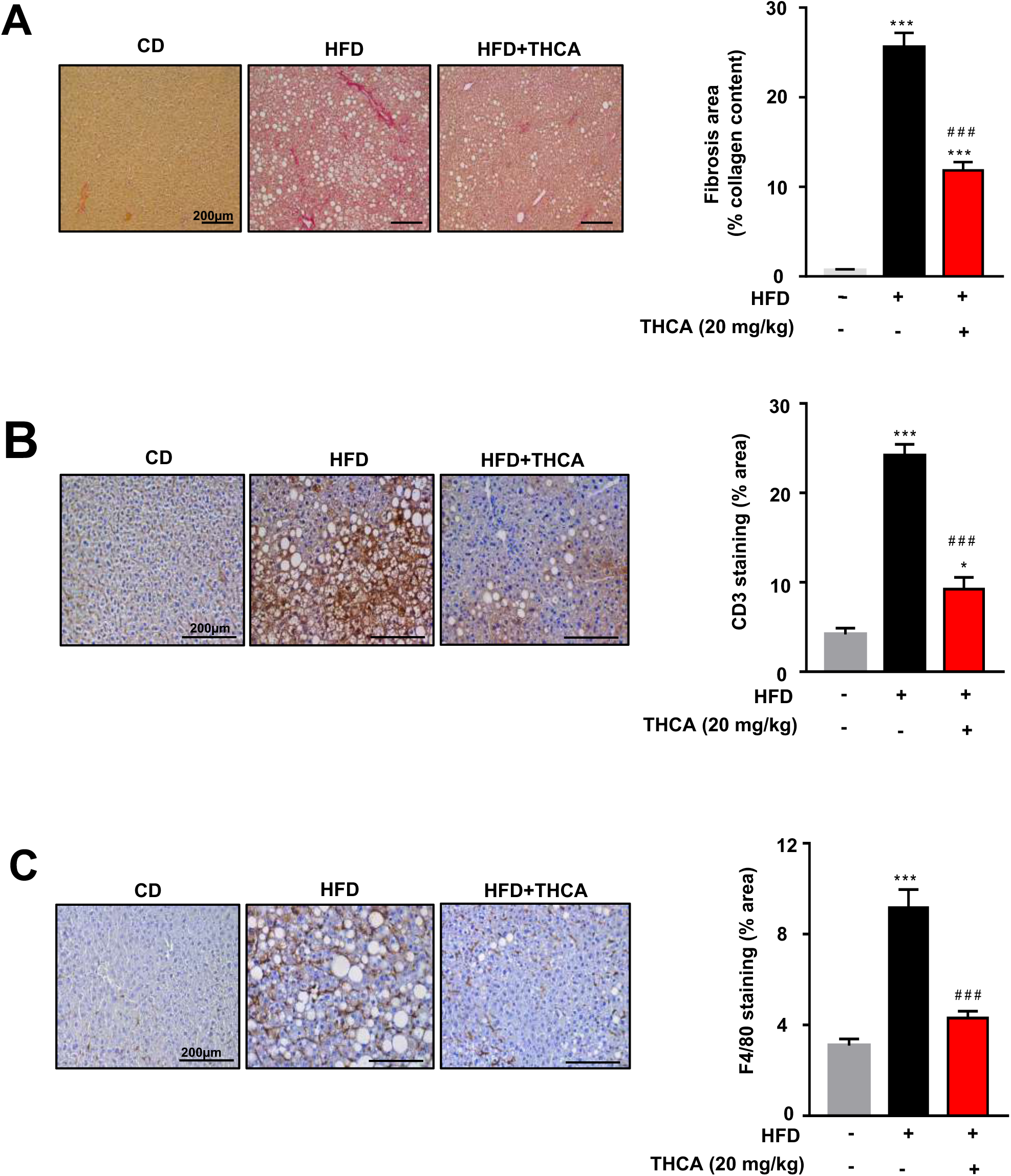
Δ^9^-THCAalleviates liver fibrosis and inflammation induced by HFD. (A) Representative images of liver collagen staining by picrosirius red dye (left panel) and the correspondingquantification (right panel) in mice fed with HFD for 23weeks and treated for the last 3 weeks with Δ^9^-THCA or vehicle compared with the corresponding CD group treated with vehicle. (B) Representative images of CD3-immunostained liver sections (left panel) and the quantification of relative CD3 positive area (right panel). (D) Representative images ofimmunostaining for F4/80 in liver sections (left panel) and the quantification (right panel). Values are expressed as mean ± SEM (n= 3-4 animals per group). *p< 0.05, ***p < 0.001 *vs*. CD group; ###p < 0.001 *vs*. HFD group (ANOVA followed by Tukey’s test).

## DISCUSSION

Salicylates and their plant sources have a long history of use in human medicine, encompassing not only the plant hormone salicylic acid and some simple derivatives like Aspirin and 5- aminosalicylic acid, but also more complex isoprenylated derivatives, like amorfrutins (Choi et al. 2015). Acidic cannabinoids are, basically, an elaborated version of isoprenylated salicylates, and, like all compounds of this class, they show a complex, multi-target biological profile, defined for Δ^9^-THCA, in terms of modulation of the transcription factor PPARγ and CB_1_R as well as in terms of inhibition of various pro-inflammatory enzymes such are PLC and COX-2 (Moreno-Sanz 2016). In previous work, we showed that Δ^9^-THCA could significantly reduce fat mass and body weight gain in a less severe murine model of HFD-induced obesity, preventing liver steatosis, adipogenesis and macrophage infiltration in fat tissues. Since obesity is closely associated to NAFLD, we wondered if THCA-A could be beneficial in this condition also in different models of induction, and, if so, what could be its molecular mechanism of action. Herein, we show that Δ^9^-THCA prevents NAFLD in mouse models of CCl_4_ and HFD-induced liver fibrosis. Importantly, the beneficial effects of Δ^9^-THCA to avert liver fibrosis persist even when the fibrogenic insults are maintained.

Chronic inflammation is the main cause of hepatic fibrogenesis, as occurs in NAFLD. In the acute phase of CCl_4_ hepatic injury, oxidative stress from damaged hepatocytes and Kupffer cells represent the initial paracrine stimulation that enhances hepatic stellate cells (HSCs) proliferation and contributes to the production of ECM, mainly collagen fibers (Sato et al. 2014). In liver fibrosis, regardless of the etiology, the initial inflammatory phase is characterized by the activation of liver-resident Kupffer cells and infiltrating immune cells that release pro-inflammatory cytokines (mainly TNFα, IFNγ and TGF-β). Hepatic inflammation triggers the activation and differentiation of HSCs from quiescent cells to myofibroblasts acquiring proliferative, pro-inflammatory and contractile properties and contributing to the production of ECM, mainly collagen fibers (Sato et al. 2014). Activation of HSCs in fibrogenesis involves distinct morphological and biochemical changes. This activation requires the coordinated changes in activity of several transcription factors. PPARγ is one such factor whose activity is decreased in activated HSCs. Indeed, there is evidence that PPARγ ligands reverse the biochemical features of HSCs activation both *in vitro* and *in vivo* (Miyahara et al. 2000; Galli et al. 2002). Since Δ^9^-THCA is a potent PPARγ ligand agonist, it is likely that it can exert an anti-fibrogenic effect by acting at different steps, firstly by reducing the number of activated HSCs and hepatic myofibroblasts, as shown by lower α-SMA^+^ cells, with a concomitant reduction of the fibrotic area in the liver and a marked suppression of the expression of fibrogenic genes, and secondly by preventing the activation of the immune system and the production of pro-inflammatory cytokines such as TNFα, IL-1β, and IL-6 (Ricote et al. 1999; Clark et al. 2000; Tontonoz and Spiegelman 2008). We found that treatment with Δ^9^-THCA greatly reduced the infiltration of macrophages into the liver in both disease models, together with a significant decrease in the expression of *Tnfa, Il1b* and *Il6*. Treatment with Δ^9^-THCA also reduced the infiltration of T cells into the liver in both models of liver fibrosis. Of note, infiltration of immune cells was markedly greater in the liver from HFD-induced NALFD mice than in the CCl_4_ model. Moreover, the fibrotic process was more diffused and accused in the DIO model than in CCl_4_-induced liver fibrosis model, since the development of fibrosis is a longer process in which fibrogenesis occurs later due to the predominance of an inflammatory action. The reduction of the immune cell infiltration and of total fibrotic area by Δ^9^-THCA treatment was more accused in the DIO NALFD model than in the CCl_4_-induced model. Our data suggest that the chronic inflammation produced in the liver is the first predominant process that originates fibrosis as a last step. In both cases, our results strongly suggest that Δ^9^-THCA accelerates tissue remodeling towards a healthy phenotype by decreasing the activation of HSCs and/or by reducing production of pro-inflammatory cytokines that, in turns, activate HSCs, subsequently reducing fibrogenesis.

CB_1_R has been shown to successfully prevent liver inflammation and fatty liver (Siegmund et al. 2007; Jeong et al. 2008; Osei-Hyiaman et al. 2008; Jourdan et al. 2017). CB_1_R is highly expressed in hepatic myofibroblasts, fully activated HSCs and in Kupffer cells, and genetic ablation of CB_1_R in Kupfer cells reduces their polarization to an M1 pro-inflammatory phenotype (Jourdan et al. 2017). In addition, pharmacological blockade of CB_1_R prevents CCl_4_-, thioacetamide- and bile duct ligation-induced liver fibrosis (Teixeira-Clerc et al. 2006; Giannone et al. 2012). Specifically, CB_1_R blockade reduces the expression of TGFβ, inhibits hepatic myofibroblasts growth and reduces accumulation of fibrogenic cells in the liver. In murine models of obesity and of alcoholic steatohepatitis, inverse agonism of CB_1_R has been shown to successfully prevent liver inflammation and fatty liver (Siegmund et al. 2007; Jeong et al. 2008; Osei-Hyiaman et al. 2008; Jourdan et al. 2017). CB_1_R is highly expressed in hepatic myofibroblasts, fully activated HSCs and in Kupffer cells. The binding of Δ^9^-THCA to cannabinoid receptors is controversial (McPartland et al. 2017), and it has been also suggested that Δ^9^-THCA can bind CB_1_R but it is not psychotropic because does not cross the brain blood barrier (Moreno-Sanz 2016). Using the same preparation of Δ^9^-THCA used in this work we have shown recently that Δ^9^-THCA binds to the orthosteric and allosteric sites of the CB_1_R and prevents collagen-induced arthritis in mice; this activity was reversed by co-treatment with SR141716, a selective CB_1_R antagonist. We described also that Δ^9^-THCA could function as a CB_1_R positive allosteric modulator (PAM) (Palomares et al. 2020b). Although we have not investigated the role of CB_1_R on the antifibrotic activity of Δ^9^-THCA, it is possible that a rapid desensitization/internalization of CB_1_R upon repeated agonist exposure could explain, in part, the beneficial effect of Δ^9^-THCA on NALFD (Rinaldi-Carmona et al. 1998; Jin et al. 1999). It could be also possible that Δ^9^-THCA could exert its positive effects in NALFD by acting as a PAM of CB_1_R. Interestingly, the anti-inflammatory lipid lipoxin, A4, is an endogenous PAM of CB_1_R (Pamplona et al. 2012), which inhibited HSCs activation (T. Zhang et al. 2020) and attenuated obesity-induced liver disease (Börgeson et al. 2015). Further research to elucidate the role of CB_1_R on the anti-fibrotic activity of Δ^9^-THCA is warranted.

Altogether, these features add to our current efforts to identify pharmacological cannabinoids with anti-inflammatory and anti-fibrotic activities, and suggest that Δ^9^-THCA, as well as non-decarboxylated *Cannabis sativa* extracts, are worth of consideration for the management of NALFD. In addition, Δ^9^-THCA could serve as a template to develop novel and chemically more stable, semi-synthetic derivatives.

## Acknowledgments

This research was funded by grants SAF2017-87701-R (EM) and BFU2017-83934-P (MTS) (Ministerio de Economía y Competitividad, Spain; co-funded with EU funds from FEDER Program); Project P12-FQM-01943 (M.T.-S.; Junta de Andalucía, Spain). CIBER Fisiopatología de la Obesidad y Nutrición is an initiative of Instituto de Salud Carlos III. This work was also partially supported by Emerald Health Biotechnology España (Cordoba, Spain). None of the funding bodies played any role in the study design, data collection and analysis, the decision to publish, or the preparation of the manuscript.

## Author contributions

BCH, IGM and FRP performed *in vitro* and in *vivo* experiments. GA isolated Δ^9^-THCA. EM, AGM and MTS managed and designed the overall study. All the authors wrote and approved the final manuscript.

## Abbreviations

ALT: Alanine aminotransferase
BW: Body weight
CB_1_R: Cannabinoid type 1 receptor
CCl_4_: Carbon tetrachloride
CD: Control diet
CLD: Chronic liver disease
CLS: Crown-like structures
DIO: Diet-induced obesity
ESLD: End-stage liver disease
GTT: Glucose tolerance test
HFD: High fat diet
HSCs: Hepatocellular stellate cells
iWAT: Inguinal white adipose tissue
LBD: Ligand binding domain
MetS: Metabolic syndrome
NAFLD: Non-alcoholic liver disease
NASH: Steatohepatitis
PPAR: Peroxisome Proliferator-Activated Receptor
PSR: Picrosirius red
TG: Triglyceride
Δ^9^-THC: Δ^9^-tetrahydrocannabinol
Δ^9^-THCA: Δ^9^-tetrahydrocannabinol acid.

---

**Table 1.**
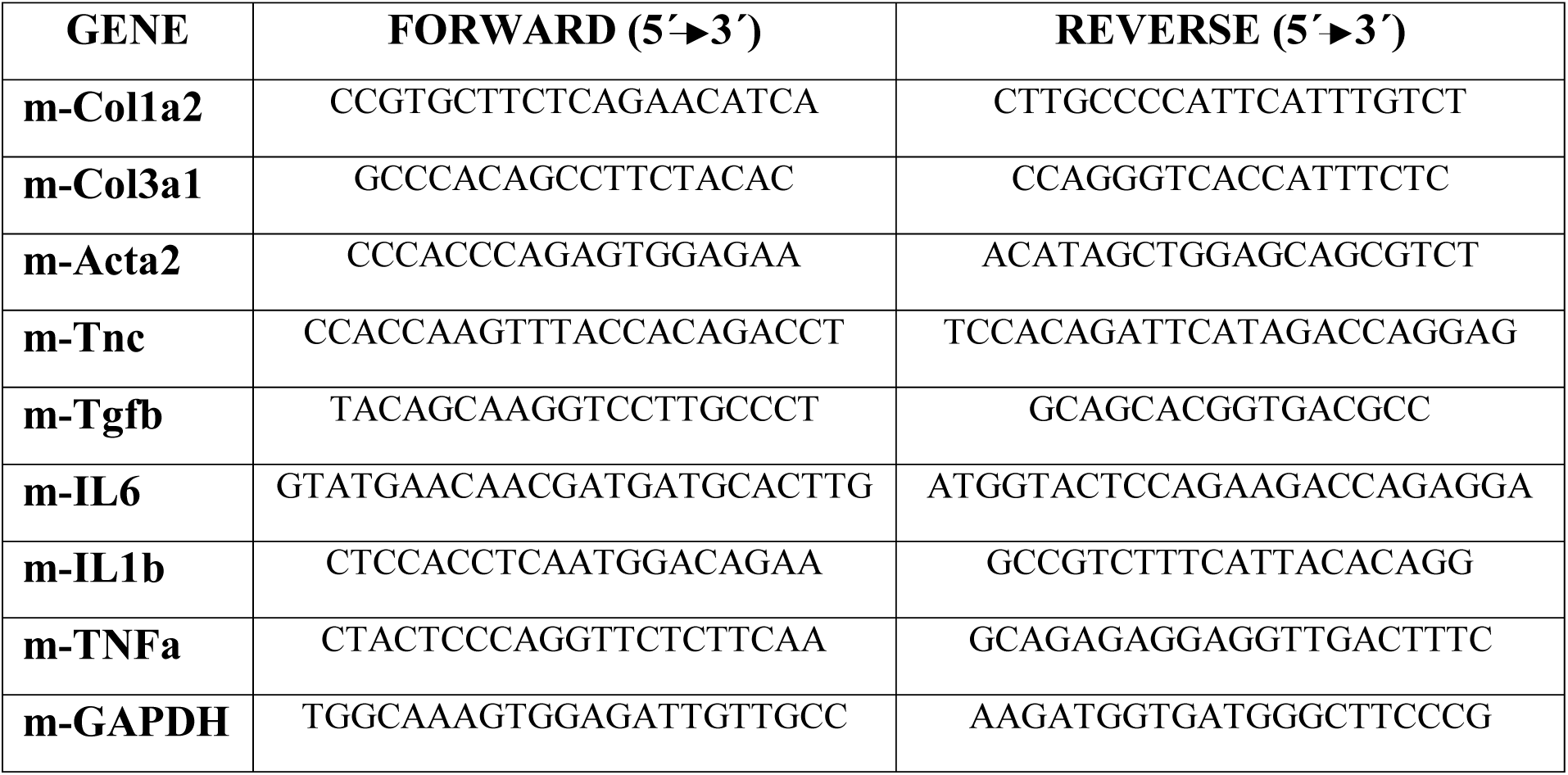
Primers used in this study

## Supplementary information

**Figure S1.**
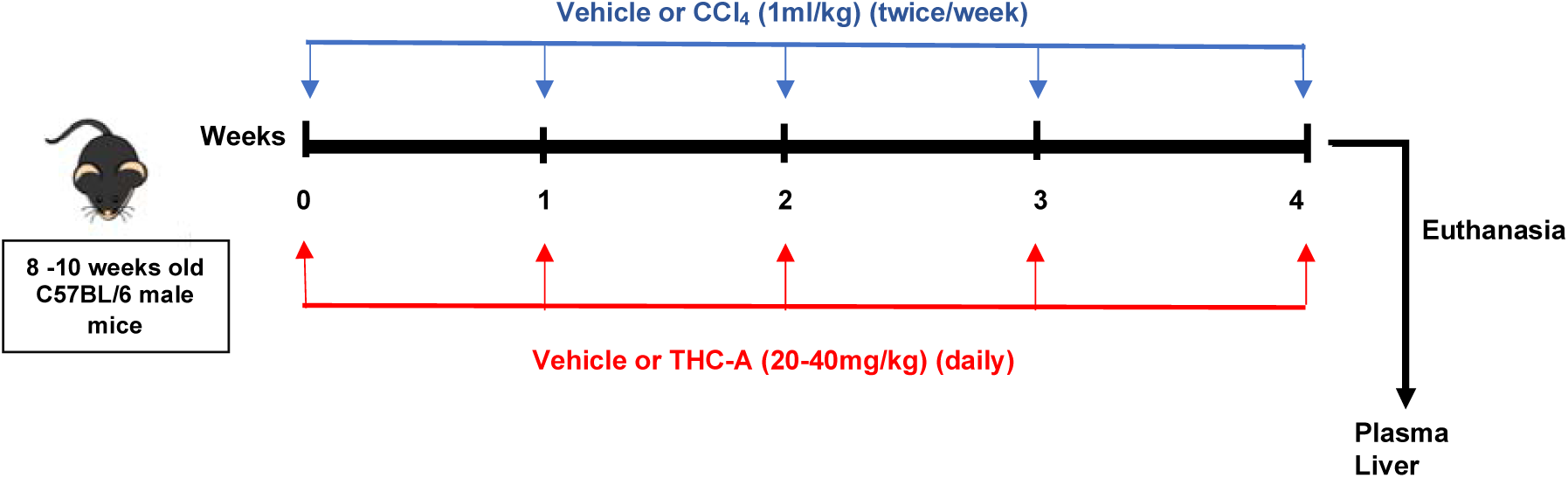
Flow-chartandtimeline ofstudydesign for CCl_4_-induced liver fibrosis model. 8-10 weeks old C57BL/6 male mice were dosed intraperitoneally (i.p.) with carbon tetrachloride (CCl_4_) at a dose of 1 ml/kg diluted in corn oil (1:4) 2 times per week for 4 weeks to induce liver fibrosis. Control mice received vehicle injections in the same way. In parallel, during the 4-week CCl_4_ treatment period, mice were injected i.p. daily with Δ^9^-THCA at doses of 20 or 40 mg/kg or with the corresponding vehicle (Ethanol: Cremophor: Saline 1:1:18). Seventy-twohours later of the last dose of CCl_4_ or vehicle, mice were euthanized, and blood and liver samples were collected for subsequent histological, metabolic and gene expression studies.

**Figure S2.**
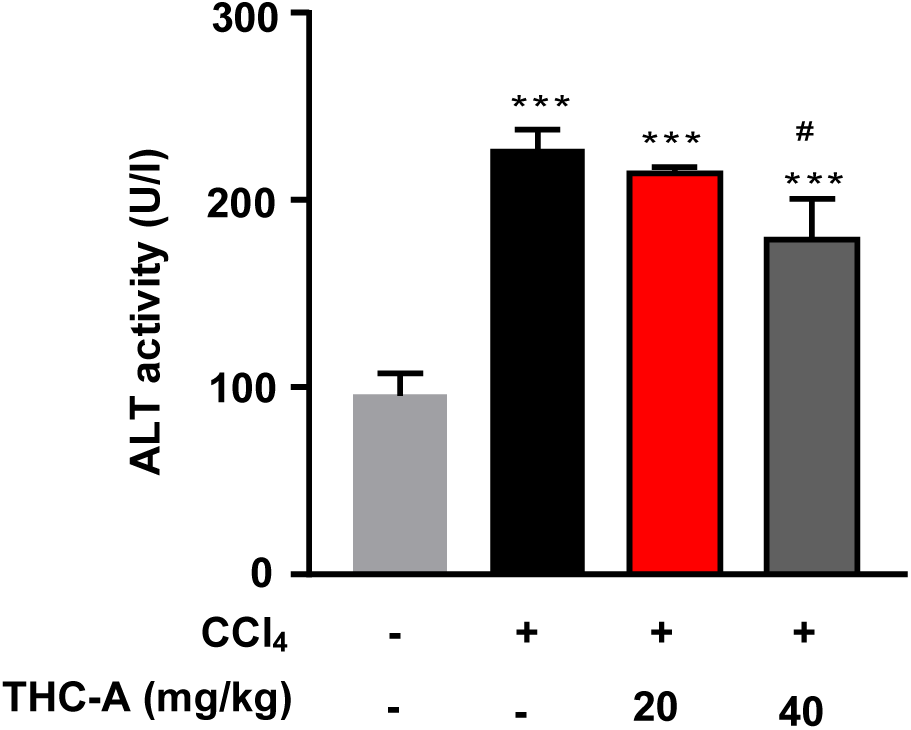
Δ^9^-THCA reduces circulating levels of serum alanine aminotransferase in liver fibrosis induced by CCl_4_. Quantification of alanine aminotransferase (ALT) activity in plasma in control, CCl_4_-vehicle and CCl_4_ + Δ^9^-THCA mice. Values are expressed as mean ± SEM (n=4-6 animals per group). ***p < 0.001 *vs*. control group; #p < 0.05 *vs*. CCl_4_group(ANOVA followed by Tukey’s test).

